# Surf4 Promotes Endoplasmic Reticulum Exit of the Lysosomal Prosaposin-Progranulin Complex

**DOI:** 10.1101/2021.03.11.435014

**Authors:** S. Devireddy, S.M. Ferguson

## Abstract

Progranulin is a lysosomal protein whose haploinsufficiency causes frontotemporal dementia while homozygous loss of progranulin causes neuronal ceroid lipofuscinosis, a lysosomal storage disease. The sensitivity of cells to progranulin deficiency raises important questions about how cells coordinate intracellular trafficking of progranulin to ensure its efficient delivery to lysosomes. In this study, we discover that progranulin interacts with prosaposin, another lysosomal protein, within the lumen of the endoplasmic reticulum (ER) and that prosaposin is required for the efficient ER exit of progranulin. Mechanistically, we identify an interaction between prosaposin and Surf4, a receptor that promotes loading of lumenal cargos into COPII coated vesicles, and establish that Surf4 is critical for the efficient export of progranulin and prosaposin from the ER. Collectively, this work demonstrates a network of interactions occurring early in the secretory pathway that promote the ER exit and subsequent lysosomal delivery of newly translated progranulin and prosaposin.

## Introduction

Heterozygous, loss-of-function mutation in the *granulin (GRN)* gene, which encodes the progranulin protein, cause a neurodegenerative disease known as frontotemporal dementia ^1–3^. Meanwhile, homozygous loss of progranulin results in neuronal ceroid lipofuscinosis, a lysosome storage disease with an earlier age of onset ^4,5^. The human progranulin protein is made up of a series of 7 repeats of the cysteine-rich granulin domain and lacks identifiable similarity to known enzymes or other proteins. The linkers between these granulin domains are cleaved within lysosomes to yield individual 6-8 kDa granulin peptides ^6–11^. Although the direct biochemical function of progranulin or the granulin fragments remains an open area of investigation, localization of progranulin and the granulins to lysosomes along with the lysosomal disease arising from *GRN* mutations indicates their important role in supporting lysosomal activity.

Given the importance of progranulin for the maintenance of normal lysosome function, mechanisms must exist to ensure the delivery of progranulin to lysosomes. Previous studies identified sortilin as a receptor for the endocytic uptake of progranulin ^12^. Prosaposin was also identified as a progranulin interacting protein that links progranulin to the cation independent mannose-6-phosphate receptor (CI-MPR) for trafficking from the trans-Golgi network to endosomes as well as to either the mannose-6-phosphate receptor or low-density lipoprotein receptor 1 (LRP1) for endocytic uptake of progranulin ^13,14^. Prosaposin has a modular domain organization that is made up of 4 saposin domains that are liberated by proteolytic cleavage within lysosomes and which function to selectively extract lipid bound molecules and present them to specific soluble enzymes within the lysosome lumen ^15,16^. Prosaposin thus functions as both a scaffold for the trafficking of progranulin as well as the precursor for lysosomal saposins. Given that *GRN*-linked frontotemporal dementia arises from progranulin haploinsufficiency, increasing the function of the progranulin encoded by the remaining wildtype copy of the *GRN* gene represents an actively investigated therapeutic strategy ^17^. However, the success of such approaches depends on detailed understanding of the mechanisms that either stimulate the activity of progranulin/granulins within lysosomes or enhance the delivery of progranulin from its site of translation at the rough endoplasmic reticulum (ER) to its site of action within the lumen of lysosomes. To address these issues, we performed quantitative imaging and biochemical analysis of progranulin traffic through the secretory pathway combined with genetic perturbations to candidate regulators of this process. In contrast to expectations that prosaposin acts solely to promote progranulin exit from the trans-Golgi network, we discovered that progranulin exit from the ER is also highly dependent on prosaposin. This observation indicated that efflux of progranulin from the ER is not a passive process and raised questions about the machinery responsible for promoting the exit of progranulin and prosaposin from the ER. Investigation of the underlying mechanism led us to identify Surf4, a sorting receptor for COPII vesicles, as a prosaposin interacting protein that is critical for the ER to Golgi trafficking of progranulin and prosaposin. Our observations support a model wherein newly translated progranulin and prosaposin interact within the lumen of the ER and bind via prosaposin to Surf4 for their packaging into COPII vesicles for delivery to the Golgi. These new findings concerning progranulin and prosaposin engaging in Surf4-dependent trafficking early in the secretory pathway complement the previous studies that defined later roles for the CI-MPR, LRP1 and sortilin at the trans-Golgi network and the plasma membrane ^12–14,18^. Each of these regulated trafficking events will contribute to how efficiently progranulin-prosaposin is delivered to lysosomes and is thus of fundamental cell biological relevance and of potential value for future strategies to enhance progranulin trafficking for therapeutic purposes in neurodegenerative diseases.

## Results

### Prosaposin is required for the efficient ER exit of progranulin

To investigate the regulatory role of prosaposin in controlling progranulin trafficking, we examined the effects of prosaposin depletion (Supp. Fig. 1A, B) on progranulin subcellular localization. In contrast to expectations arising from the proposed role for prosaposin in promoting CI-MPR-dependent sorting of progranulin at the TGN ^14^, we observed that while progranulin localized to LAMP1 labeled lysosomes in control (Fig. 1A), progranulin was absent from lysosomes in prosaposin depleted cells (Fig. 1B, Supp. Fig. 1C). Although a portion of the progranulin localized to the Golgi in the prosaposin depleted cells, we were struck by the fact that most of the progranulin was not in the Golgi and instead was in the ER (Fig. 1B, Supp. Fig. 1C). In contrast (and consistent with a previous publication ^14^), prosaposin still localized to lysosomes in progranulin knockout cells (Supp. Fig. 1D). These results also provided a robust validation of progranulin immunofluorescence specificity.

**Figure 1:**
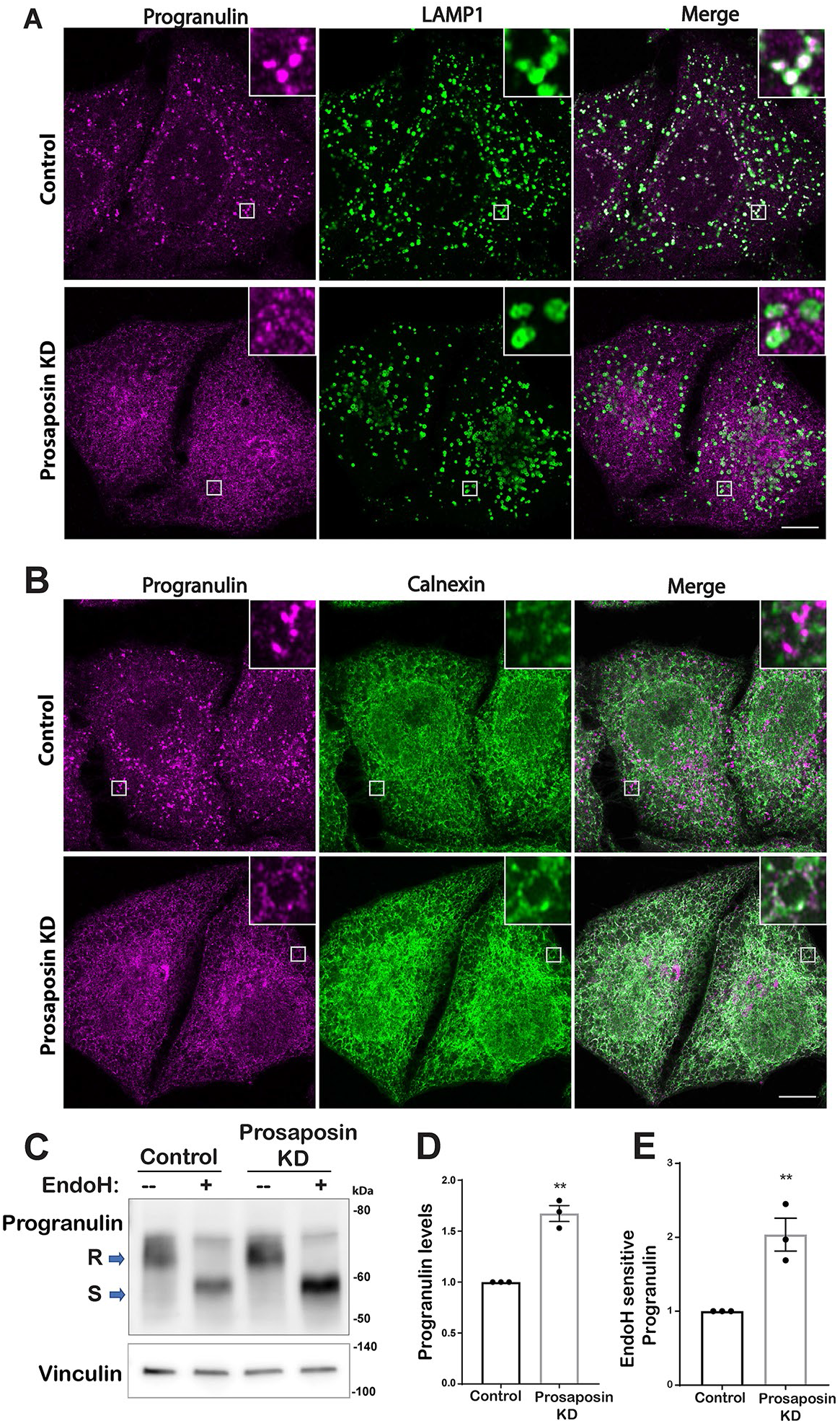
Prosaposin is required for the efficient ER exit of progranulin. (A, B) Confocal immunofluorescence images show progranulin subcellular localization together with LAMP1 labeled lysosomes (A), and the ER marker (Calnexin) (B) in control and prosaposin knockdown (KD) HeLa cells. Scale Bars, 10µm. (C) Immunoblot analysis of cell lysates from control siRNA and prosaposin siRNA transfections without (--) and with (+) endoglycosidase H (Endo H) enzyme treatment. S indicates Endo H sensitive, R - Endo H resistant form. Vinculin was used as a loading control. (D, E) Quantification of total progranulin protein levels (D) and the Endo H sensitive form of progranulin (E), normalized to vinculin in cells treated with control versus and prosaposin siRNAs (n=3; mean ± SEM; unpaired *t* test; \*\**p* value <0.01).

To independently assess the levels of ER-localized progranulin, we tested for sensitivity to endoglycosidase H (Endo H), a bacterial enzyme that selectively recognizes and deglycosylates proteins with the high mannose and hybrid N-linked glycans found on newly translated proteins within the ER while the glycans on proteins that have passed through the cis-Golgi become resistant to Endo H ^19^. We found that the majority of full length progranulin is in the Endo H sensitive, ER resident form (Supp. Fig. 2A, B). To further validate the conclusions of this assay, we analyzed the media samples for secreted progranulin and found only the Endo H resistant form (Supp. Fig. 2A). This important control confirms progranulin robustly acquires the expected Endo H resistance as it traverses the secretory pathway. Consistent with impaired progranulin delivery to lysosomes (where it is proteolytically processed into granulins ^7^), prosaposin depletion was accompanied by an increase in total progranulin levels (Fig. 1C, D). This increase was paralleled by an increase in the Endo H sensitive fraction but with no significant change in Endo H resistant fraction (Fig. 1C, E). Thus, both imaging and Endo H resistance assays demonstrated that prosaposin is required for the efficient ER-to-Golgi trafficking of progranulin.

**Figure 2:**
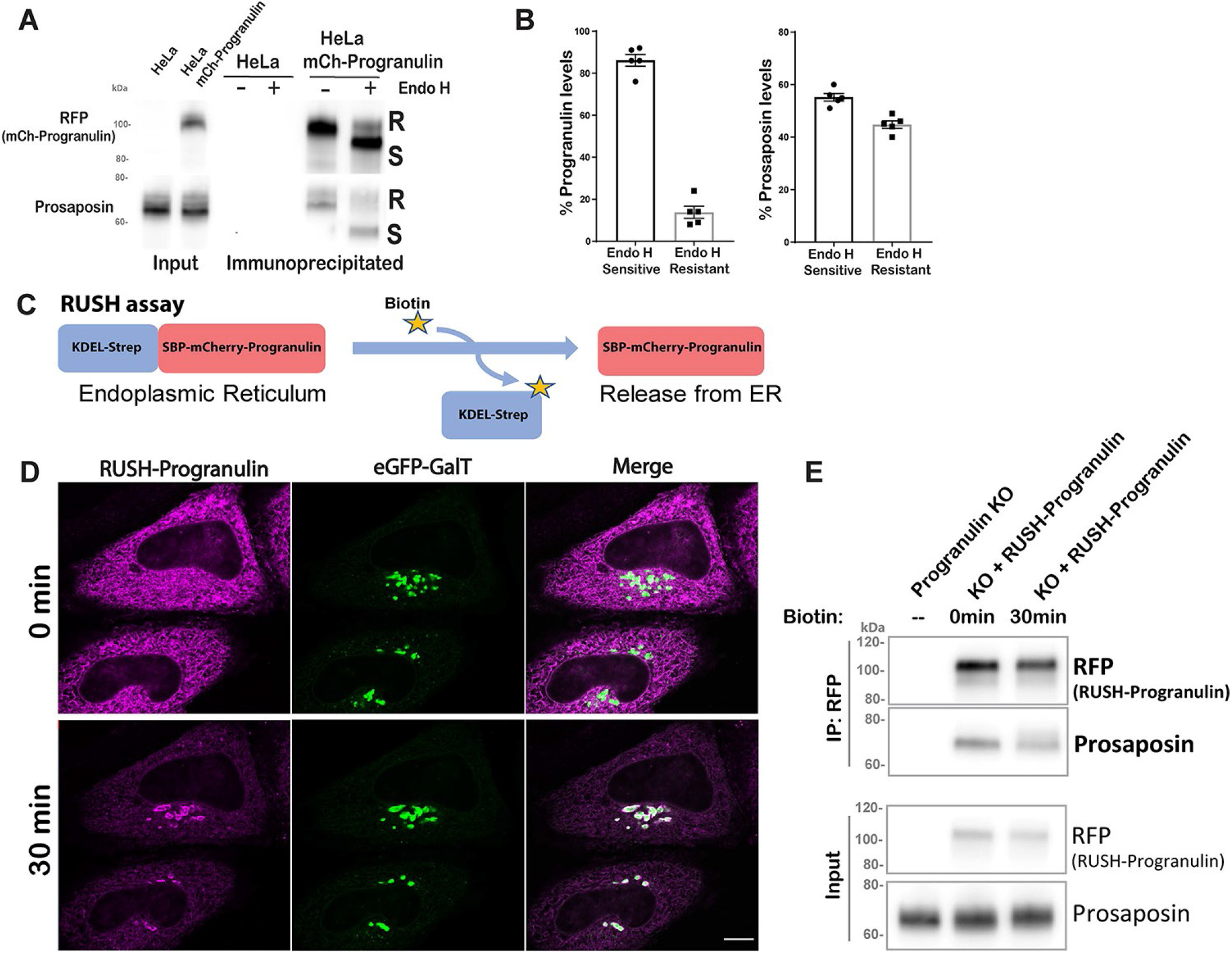
Progranulin interactions with prosaposin begin in the ER. (A) Immunoprecipitates from untransfected HeLa cells and HeLa cells stably expressing mCherry-Progranulin were treated with and without Endo H and analyzed by immunoblotting. (B) Quantification of the percentages of Endo H sensitive and resistant forms in the Progranulin-Prosaposin complex (n=5 independent experiments; mean ± SEM). (C) Schematic of RUSH (Retention using selective hooks) strategy: mCherry-Progranulin fused to streptavidin binding peptide (SBP) is retained in the ER via its interaction with KDEL-streptavidin. Addition of biotin disrupts this interaction and causes the synchronous release of SBP-mCherry-Progranulin from the ER. (D) HeLa cells were co-transfected with RUSH-progranulin and eGFP-GalT (Golgi marker) one day before imaging. Live cell confocal images show the localization of RUSH-progranulin at 0min and 30min of biotin addition (scale Bar, 10µm). (E) Immunoprecipitation of RUSH-progranulin using RFP-Trap beads was performed at 0min and at 30min of biotin treatment and immunoblots were probed for progranulin and prosaposin proteins. Ribosomal S6 was used as a loading control. Similar results were observed in three independent experiments.

### Prosaposin interactions with progranulin begin in the ER

As prosaposin depletion causes the buildup of progranulin in ER, we hypothesized that progranulin interactions with prosaposin must be initiated early in the secretory pathway. To identify the subcellular location of such interactions, Endo H sensitivity assays were performed on the immunoprecipitated complexes from cells that stably express mCherry tagged-progranulin at near endogenous levels and they revealed a robust presence of Endo H sensitive (ER form) prosaposin in complex with progranulin (Fig. 2A, B).

To further test for progranulin-prosaposin interaction within the ER, we employed a combination of the RUSH (Retention Using Selective Hooks) strategy ^20^ along with immunoprecipitation. To this end, we generated an mCherry tagged-progranulin reporter protein fused to streptavidin binding peptide (SBP-mCherry-progranulin, referred as RUSH-progranulin) and co-expressed it with KDEL-Streptavidin in order to achieve biotin regulated retention of RUSH-progranulin in the ER (Fig. 2C). Prior to the addition of biotin, RUSH-progranulin was retained in the ER compartment (Fig. 2D). Following the addition of biotin, RUSH-progranulin was released from the ER and accumulated within the Golgi compartment with a peak near 30 minutes (Fig. 2D) before eventually reaching the lysosomes (Supp. Fig. 5B). We next evaluated the RUSH-progranulin interactions with endogenous prosaposin without biotin and 30 minutes after adding biotin and found comparable levels of prosaposin interaction for both the ER and Golgi localized pools of progranulin (Fig. 2E). Collectively, these results establish that the prosaposin-progranulin interaction begins in the ER compartment and that the formation of this protein complex facilitates the ER exit of progranulin.

**Figure 3:**
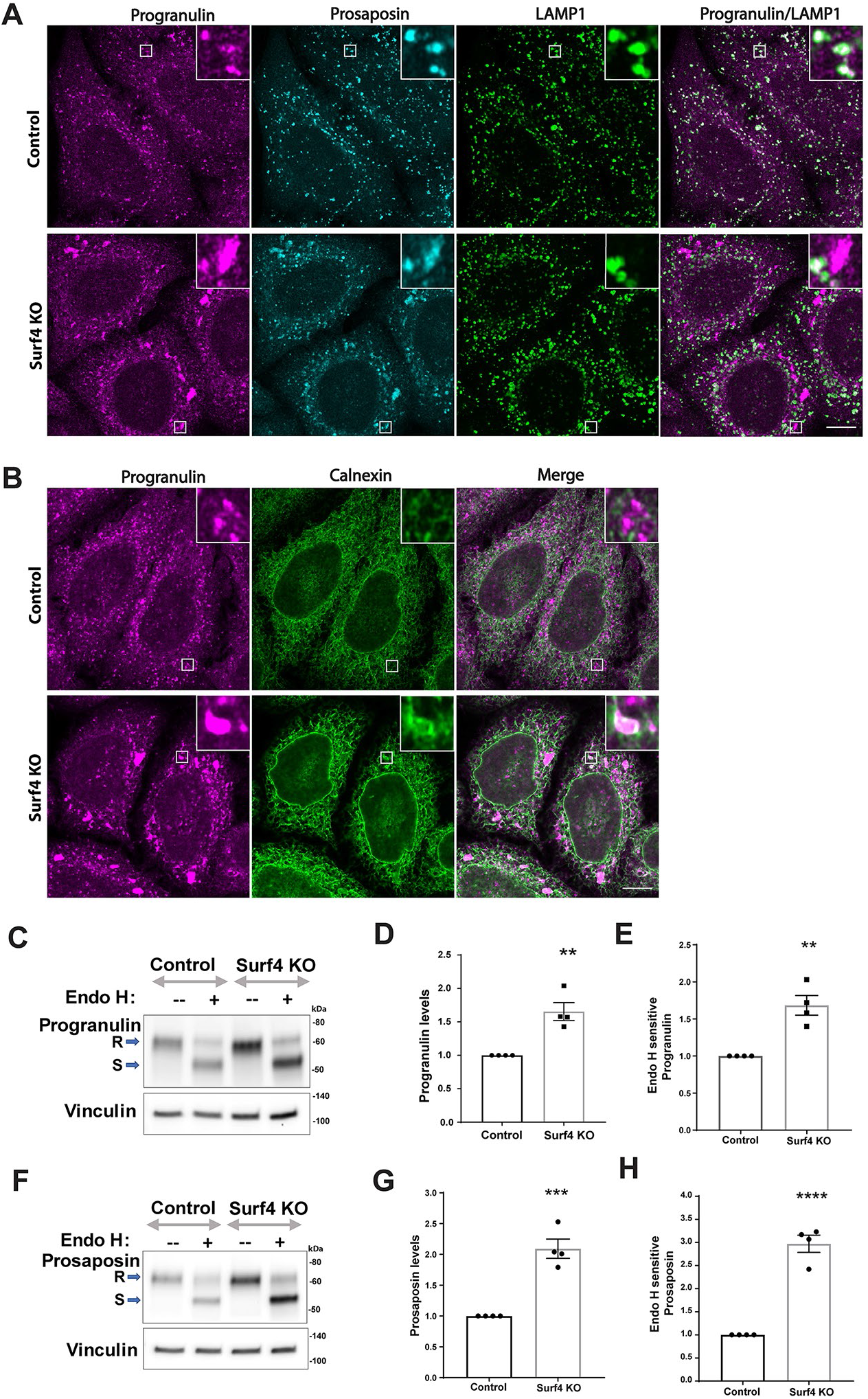
Surf4 is required for the trafficking of both progranulin and prosaposin. (A) Confocal immunofluorescence analysis of progranulin, prosaposin together and LAMP1 in control and Surf4 KO cells. (B) Confocal immunofluorescence images of progranulin and calnexin, an ER marker in control and Surf4 KO cells. Scale Bars, 10 µm. (C, F) Immunoblot analysis of cell lysates treated without (--) and with (+) Endo H shows both total levels of the respective proteins and their Endo H sensitive versus resistant forms. S denotes Endo H sensitive and R indicates Endo H resistant. Vinculin was used as a loading control. Quantification of immunoblots shows an increase in the levels of total progranulin (D) and prosaposin (G) as well as their Endo H sensitive forms (E, H) in Surf4 KO cells (n=4 independent experiments; mean ± SEM; unpaired *t* test; \*\**p*<0.01; \*\*\**p*<0.001; \*\*\*\**p*<0.0001).

**Figure 4:**
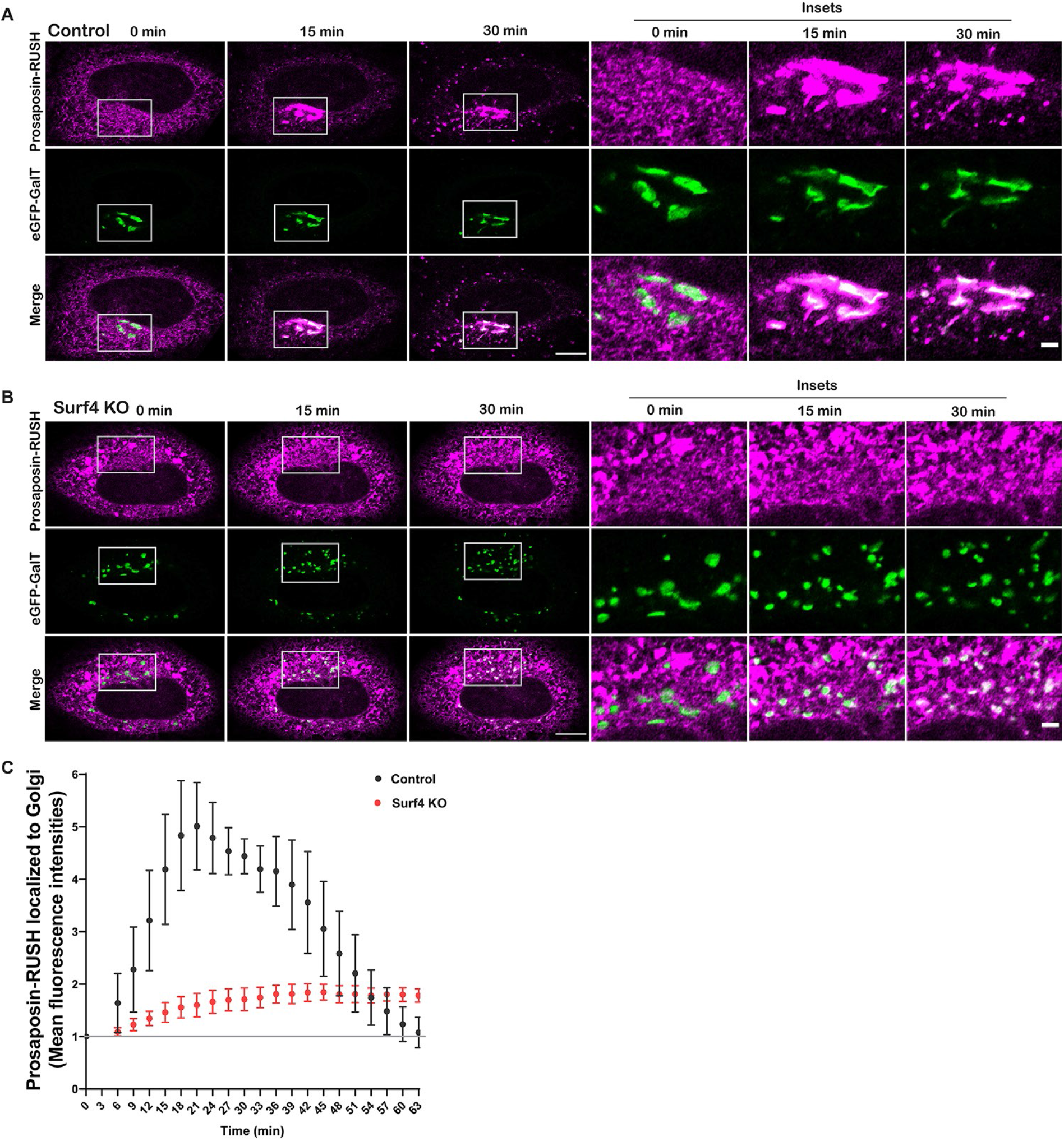
Surf4 is required for the efficient exit of prosaposin from the ER. (A, B) Live cell confocal images of prosaposin-RUSH traffic from ER are shown for control and Surf4 KO cells at the indicated timepoints after biotin addition. Magnified insets show the localization of prosaposin to eGFP-GalT labeled Golgi at the indicated times of biotin treatment. Scale Bars, 10µm. Insets, 2 µm. (C) The graphs show the kinetics of prosaposin-RUSH delivery to the Golgi in control and Surf4 KO cells. The mean fluorescence intensities of Prosaposin-RUSH localized to Golgi compartment after biotin addition were normalized to the mean fluorescence intensities before biotin (baseline marked by gray line) and plotted at the indicated times of biotin treatment. Data were collected from four independent experiments with n= 9-14 cells per experiment; Error bars show mean ± SEM.

**Figure 5:**
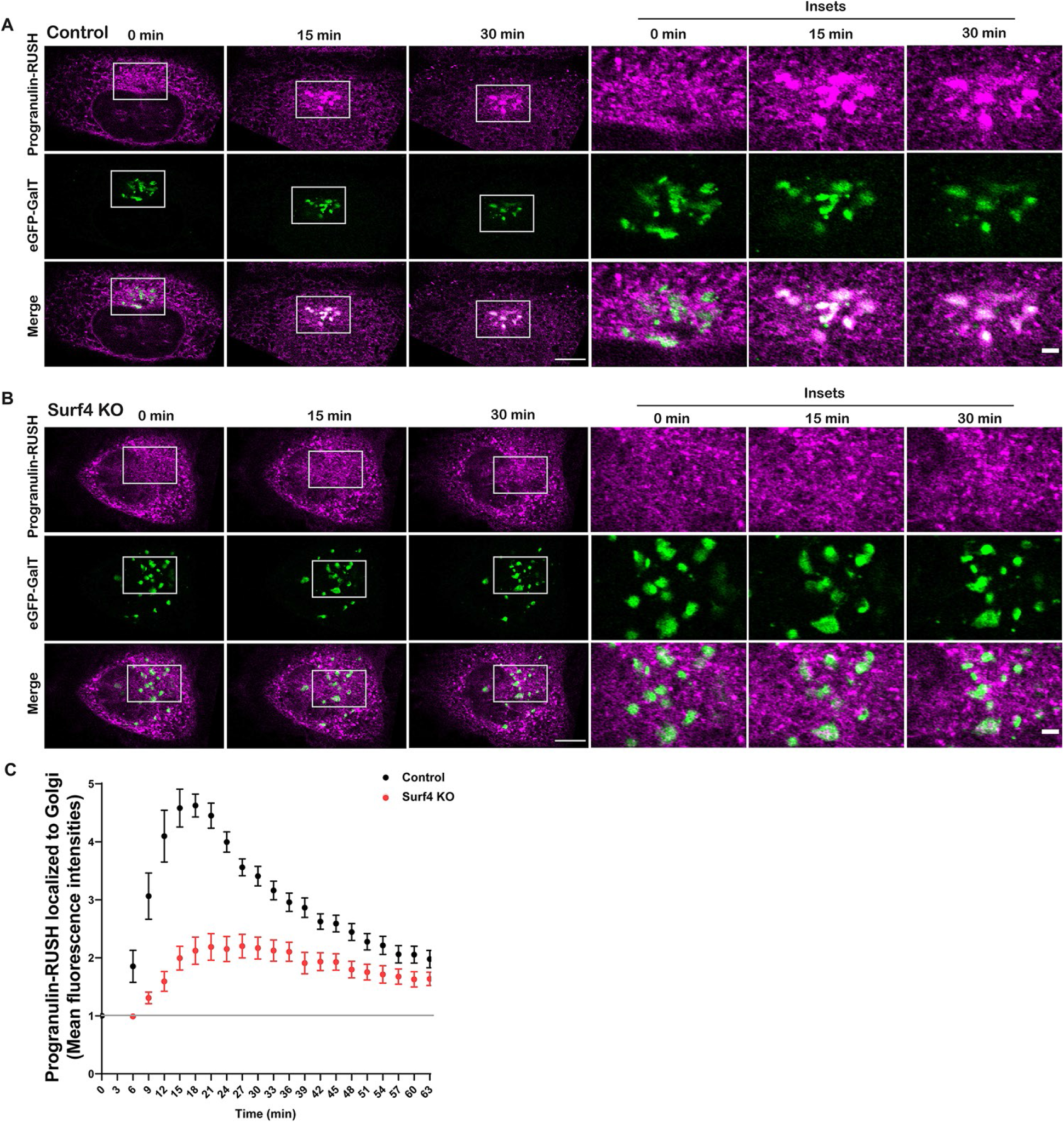
Surf4 is required for the efficient exit of progranulin from the ER. (A, B) Live cell confocal images show Progranulin-RUSH traffic from ER in control and Surf4 KO cells at the indicated timepoints after biotin addition. Magnified insets show the localization of progranulin to eGFP-GalT labeled Golgi at the indicated times of biotin treatment. Scale Bars, 10µm. Insets, 2 µm. (C) The graph shows the kinetics of Progranulin-RUSH accumulation in the Golgi in control and Surf4 KO cells. The mean fluorescence intensities of Progranulin-RUSH localized to Golgi after biotin addition were normalized to the mean fluorescence intensities before biotin (baseline marked by gray line) and plotted at the indicated times of biotin treatment. Data were collected from 10 cells; Error bars show mean ± SEM.

### Surf4 is required for trafficking of progranulin and prosaposin out of the ER

A significant fraction of newly made proteins are thought to exit the ER via a process of “bulk flow” wherein they are non-selectively captured within the fluid phase of COPII vesicles that bud from the ER and deliver their contents to the cis-Golgi ^21^. However, the dependence of progranulin on prosaposin for its ER exit argued against a passive process of bulk flow and instead suggested the need for binding to a receptor to promote their trafficking out of the ER. In contrast to bulk flow, a subset of ER lumenal proteins are selectively recruited to budding COPII vesicles via transmembrane cargo receptors ^22^. So far, only a few ER cargo receptors have been identified. Of these, ERGIC-53 and Surf4 are best characterized for their ability to promote the ER export of lumenal proteins. More recently CLN6 was shown to recruit some lysosomal enzymes and present them to CLN8 for ER export ^23,24^. However, we ruled out a requirement for CLN6/CLN8 in the trafficking of progranulin based on our observation of normal lysosome localization of progranulin in fibroblasts from the *nclf* mutant mouse that lacks CLN6 (Supp. Fig. 3). Although progranulin is a glycoprotein and thus a candidate for binding to ERGIC-53 which recognizes cargoes based on their glycosylation, progranulin’s dependence on prosaposin for ER exit, argued against direct binding of progranulin to ERGIC-53 as a major route of ER exit. We therefore focused on Surf4, a receptor for a growing list of ER cargoes ^25–28^.

To test the role of Surf4, we generated Surf4 KO cells by CRISPR/Cas9 genome editing (Supp. Fig. 4A). In Surf4 KO cells, progranulin and prosaposin were no longer enriched in lysosomes and instead built up within the ER (Fig. 3A, B). This ER localization included multiple highly concentrated foci (Fig. 3A, B). We further evaluated the ER localization of progranulin and prosaposin by Endo H sensitivity assays that revealed an increased abundance of the Endo H-sensitive (ER resident) forms of both progranulin and prosaposin in the Surf4 KO cells (Fig. 3C-H). In contrast, although cathepsin D protein levels were also slightly elevated in the Surf4 KO cells (possibly reflecting changes in lysosome homeostasis arising from progranulin and prosaposin delivery defects), cathepsin D localization to lysosomes was unaffected (Supp. Fig. 4B-D). Altogether, these experiments show that Surf4 is selectively required for the efficient delivery of progranulin and prosaposin to lysosomes and point to a key role for Surf4 in their ER export.

### Surf4 is required for the efficient ER exit of prosaposin and progranulin

Steady-state localization of prosaposin and progranulin shows that these proteins are building up in ER in the absence of Surf4. However, calculating the extent of cargo trafficking defects at steady state is obscured by continuous fractional transfer of cargo. To directly test the effects of Surf4 loss on the ER to Golgi trafficking kinetics of prosaposin and progranulin, we employed the RUSH strategy to allow temporally controlled release of fluorescently labelled cargo from ER combined with live cell imaging ^20^. As expected, prosaposin-RUSH was retained in the ER before the addition of biotin (Fig. 4A) and was delivered to LAMP1-GFP labeled lysosomes in the presence of biotin (Supp. Fig. 5A). In control cells, prosaposin-RUSH reached the Golgi compartment within ∼20min after ER release while in Surf4 KO cells, prosaposin-RUSH remained in the ER and failed to enrich at the Golgi following biotin addition (Fig. 4A-C, Supp movie 1 and 2). Progranulin-RUSH also showed the expected biotin-regulated trafficking to lysosomes and was highly dependent on Surf4 for its ER to Golgi trafficking in Surf4 KO cells (Fig. 5A-C, Supp movie 3 and 4, Supp. fig. 5B). Collectively, these quantitative analyses of the dynamic trafficking of prosaposin and progranulin establish a critical role for Surf4 in their delivery from the ER to the Golgi. Notably, ∼6.6 fold reduction in the ER to Golgi traffic of prosaposin-RUSH in Surf4 KO cells stands out compared to what has been reported for other Surf4-dependent cargos and CLN6/CLN8 dependent lysosomal hydrolase cargos ^23,25–27^. This demonstrates the major extent to which Surf4 prioritizes the ER export of prosaposin (and by extension progranulin).

### Surf4 binds prosaposin in the ER and facilitates the ER exit of progranulin-prosaposin complex

Consistent with its role in controlling traffic between the ER and cis-Golgi ^25,29^, GFP-Surf4 localized predominantly to the ER and its expression rescued the large punctate ER accumulation phenotype of progranulin in Surf4 KO cells (Fig. 6A). The RUSH assays suggested that Surf4 acts as a receptor to mediate ER exit of prosaposin and progranulin. However, we also observed fragmentation of the Golgi in Surf4 KO cells (eGFP-GalT images in Fig. 4B, 5B), consistent with the previous reports indicating the role of Surf4 in the maintenance of Golgi morphology ^29^. A direct role for Surf4 as a receptor for the packaging of the progranulin-prosaposin complex into COPII vesicles requires an interaction between prosaposin and Surf4. We tested this through immunoprecipitation experiments and found that prosaposin and progranulin selectively co-purified with Surf4 (Fig. 6B). In addition, prosaposin over-expression enhanced the interaction between progranulin (endogenous) and Surf4 (Fig. 6B). Furthermore, GFP-Surf4 expression completely rescued the lysosomal localization of progranulin (Fig. 6C, D) and prosaposin (Fig. 6E, F) in Surf4 KO cells. These results establish the specificity of the Surf4 KO impact on progranulin and prosaposin trafficking and further support a model wherein prosaposin promotes the ER export of progranulin by acting as a linker between progranulin and Surf4.

**Figure 6:**
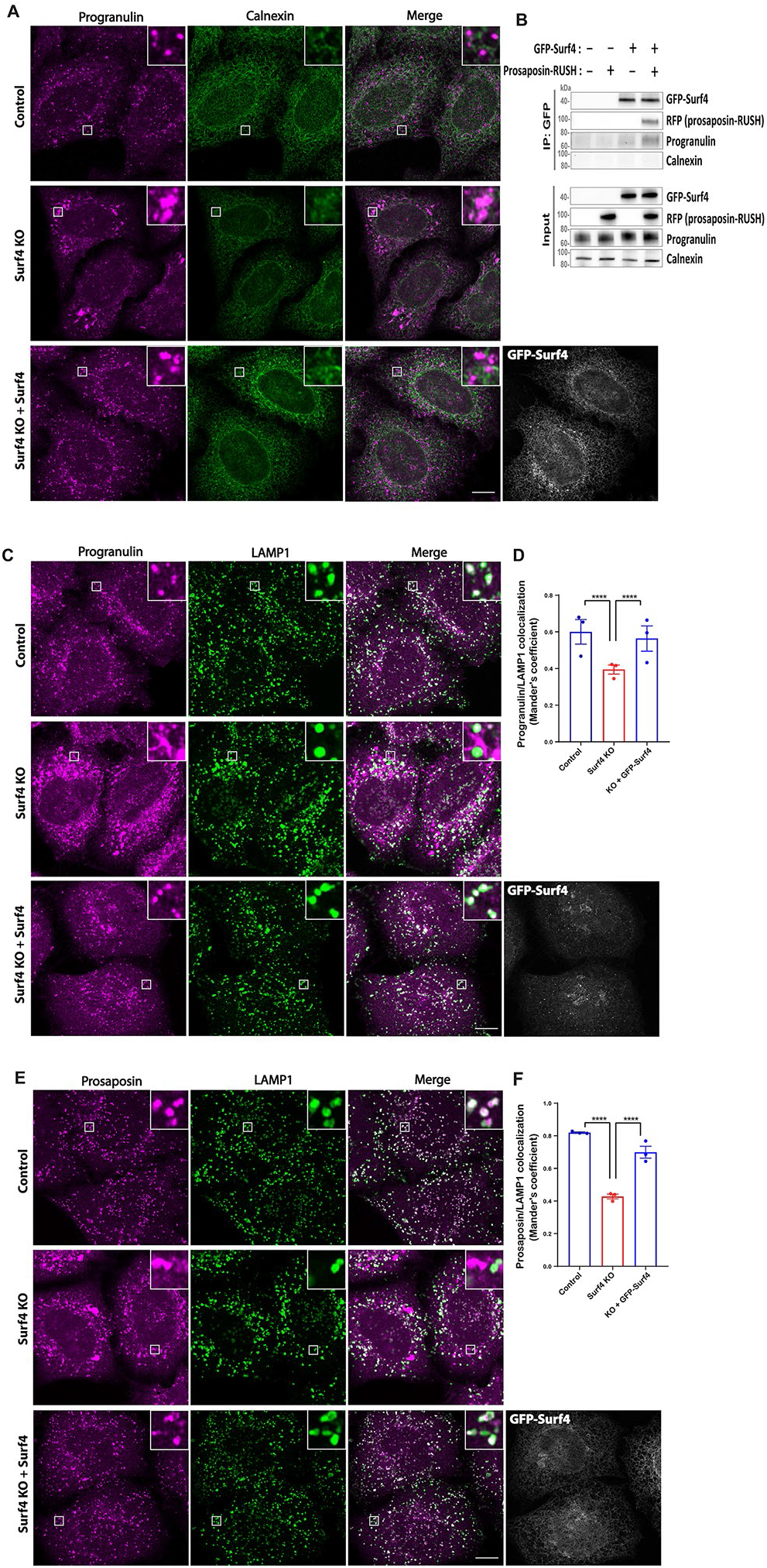
Surf4 binds prosaposin-progranulin protein complex in the ER and facilitates its ER exit. (A) Immunofluorescence images show progranulin together with calnexin (ER protein) in control, Surf4 KO and Surf4 KO cells expressing GFP-Surf4. (B) HeLa cell lysates transfected with prosaposin-RUSH and GFP-Surf4 alone or together were subjected to immunoprecipitation using GFP-Trap beads. To capture the ER localized interactions, prosaposin-RUSH was retained in the ER without the addition of biotin. Immunoblot analysis of the immunoprecipitated fractions detected prosaposin-RUSH only when expressed with GFP-Surf4 indicating the interaction between prosaposin and Surf4. Progranulin (endogenous) was detected at a comparatively higher level in immunoprecipitated fractions of prosaposin-RUSH and GFP-Surf4 than GFP-Surf4 alone suggesting that prosaposin acts as a linker to mediate progranulin interactions with Surf4. Absence of calnexin in immunoprecipitated fractions serves as a control to show the specificity of Surf4 interactions with prosaposin-progranulin complex. Similar results were observed in three independent experiments. (C, E) Immunofluorescence analysis of progranulin, prosaposin and LAMP1 in control, Surf4 KO and Surf4 KO cells expressing GFP-Surf4 (scale Bars, 10µm). (D, F) Image quantification shows the colocalization coefficients of progranulin and prosaposin with LAMP1. Data were collected from three independent experiments with n= 20 cells quantified per experiment. Error bars show mean ± SEM. One-way ANOVA with Bonferroni’s multiple comparisons test, \*\*\*\**p*<0.0001.

### A C-terminal motif is critical for Surf4-dependent ER exit of progranulin and prosaposin

In order for Surf4 to promote progranulin-prosaposin efflux from the ER via COPII vesicles, Surf4 must interact with the COPII coat via cytoplasmic sorting signals ^22,30^. We therefore next sought to identify and mutate putative sorting motifs within the cytoplasmic region of Surf4 and determine the impact of such perturbations on the ability of Surf4 to support the trafficking of prosaposin and progranulin. Erv29p, the budding yeast homolog of Surf4, was experimentally determined to possess four transmembrane domains with both N- and C-termini localized to cytoplasm ^31^. Although the C-terminus of Surf4 lacks a canonical COPII interacting motif such as a pair of phenylalanines at the very C-terminus such as those found in other cargo receptors such as ERGIC-53/LMAN1 and p24 proteins ^30,32,33^, a striking feature arising from our analysis of the Surf4 cytoplasmic C-terminus was the presence of a stretch 20 amino acids that are highly conserved across multiple distantly related species (Fig. 7A). To functionally test the role for this motif in Surf4-dependent traffic of progranulin-prosaposin, we generated GFP-Surf4 with a 15 amino acid deletion within this highly conserved region (Surf4Δmotif) and found that it failed to rescue the ER accumulation of progranulin and prosaposin in Surf4 KO cells (Fig. 7B-E). We also generated a GFP-Surf4 construct wherein a pair of adjacent phenylalanines within the conserved motif was changed to alanines (GFP-Surf4 FF→AA) and found it also failed to rescue the ER accumulation of progranulin and prosaposin (Fig. 7B-E). Together, these results indicate that the ability of Surf4 to support ER export is dependent on a conserved motif within its cytoplasmic C-terminus. We speculate that this region might play a role in mediating COPII interactions and suggest that biochemical and structural efforts will be required in the future to test such a model.

**Figure 7:**
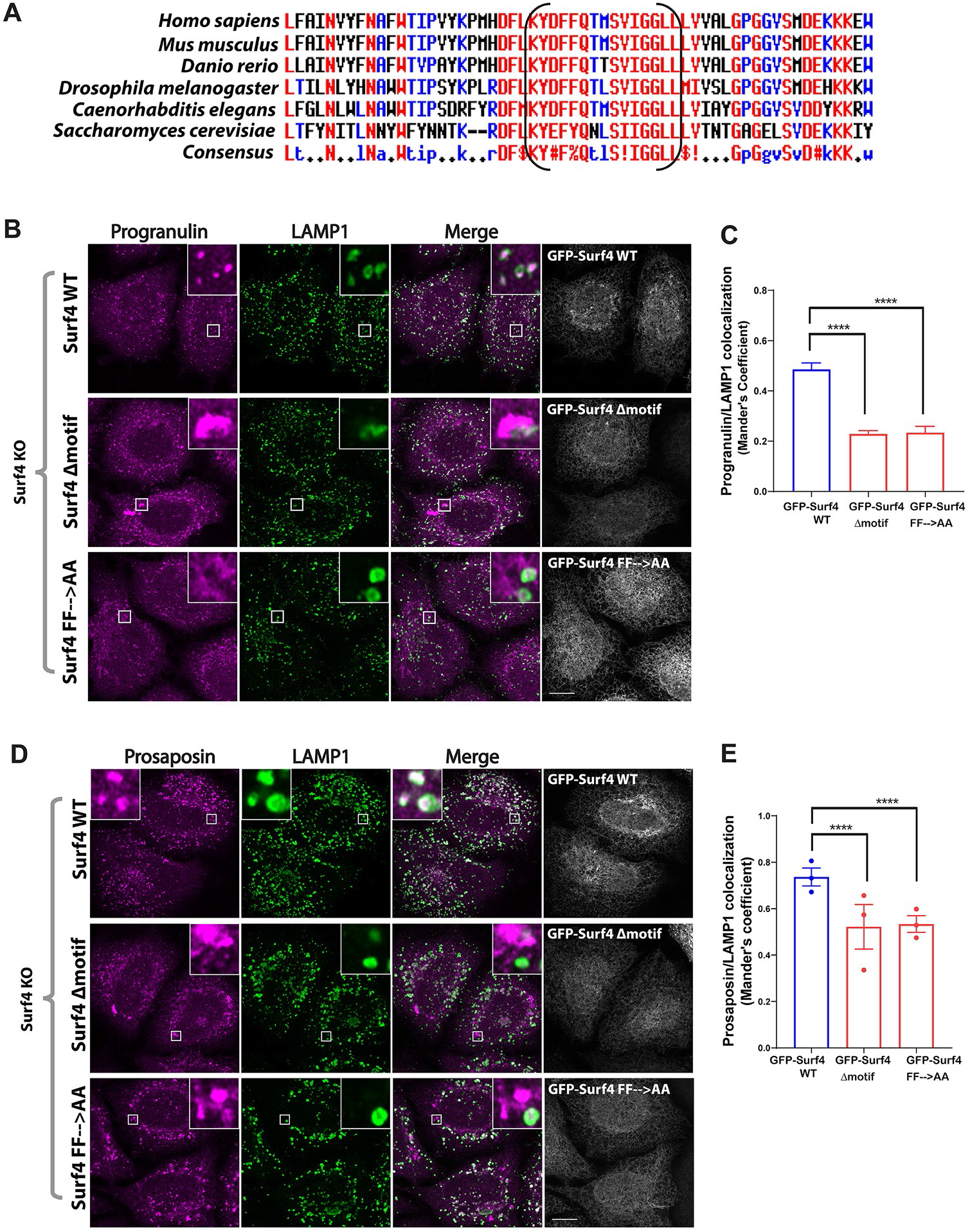
Identification of a cytoplasmic motif that is critical for Surf4-dependent trafficking of progranulin and prosaposin. (A) Analysis of Surf4 proteins across multiple species revealed a highly conserved stretch of 15 amino acids within the cytoplasmic C-terminus. Brackets highlight the region that was subsequently deleted in the GFP-Surf4Δmotif mutant. Amino acids with high consensus are shown in red color font, low consensus in blue font, and neutral in black font by MultAlin. (B, D) Immunofluorescence images show progranulin, prosaposin and LAMP1 localization in Surf4 KO cells expressing GFP-Surf4 WT, GFP-Surf4Δmotif and GFP-Surf4 FF→AA (mutation of the diphenylalanine within the conserved motif). Scale Bars, 10µm. (C) Image quantification shows the colocalization coefficients of progranulin with LAMP1. Data were collected from one experiment with 20 cells. Error bars show mean ± SEM. Kruskal-Wallis test with Dunn’s multiple comparisons test, \*\*\*\**p*<0.0001. (E) Image quantification shows the colocalization coefficients of prosaposin with LAMP1 and data were collected from three independent experiments with 20 cells quantified per condition per experiment. Error bars show mean ± SEM. Kruskal-Wallis test with Dunn’s multiple comparisons test, \*\*\*\**p*<0.0001.

## Discussion

Protein sorting within successive compartments in the secretory pathway is essential for ensuring the degradative activity of lysosomes by controlling the delivery of hydrolases and their regulatory factors ^34^. It has long been known that an important regulated step supporting the delivery of lysosome lumenal proteins takes place at the trans-Golgi network via their interactions with proteins such as mannose-6-phosphate receptors that promote sorting into the endolysosomal pathway ^35^. However, in this study, we identified a critical role for an interaction between prosaposin and progranulin in promoting the export of progranulin from the ER. We furthermore identified Surf4, a receptor which recruits lumenal cargos into COPII vesicles as a major mediator of ER export of the progranulin-prosaposin complex. These discoveries demonstrate how a series of protein-protein interactions within the ER prioritizes the trafficking of progranulin and prosaposin and has a major impact on their delivery to lysosomes (Figure 8).

**Figure 8:**
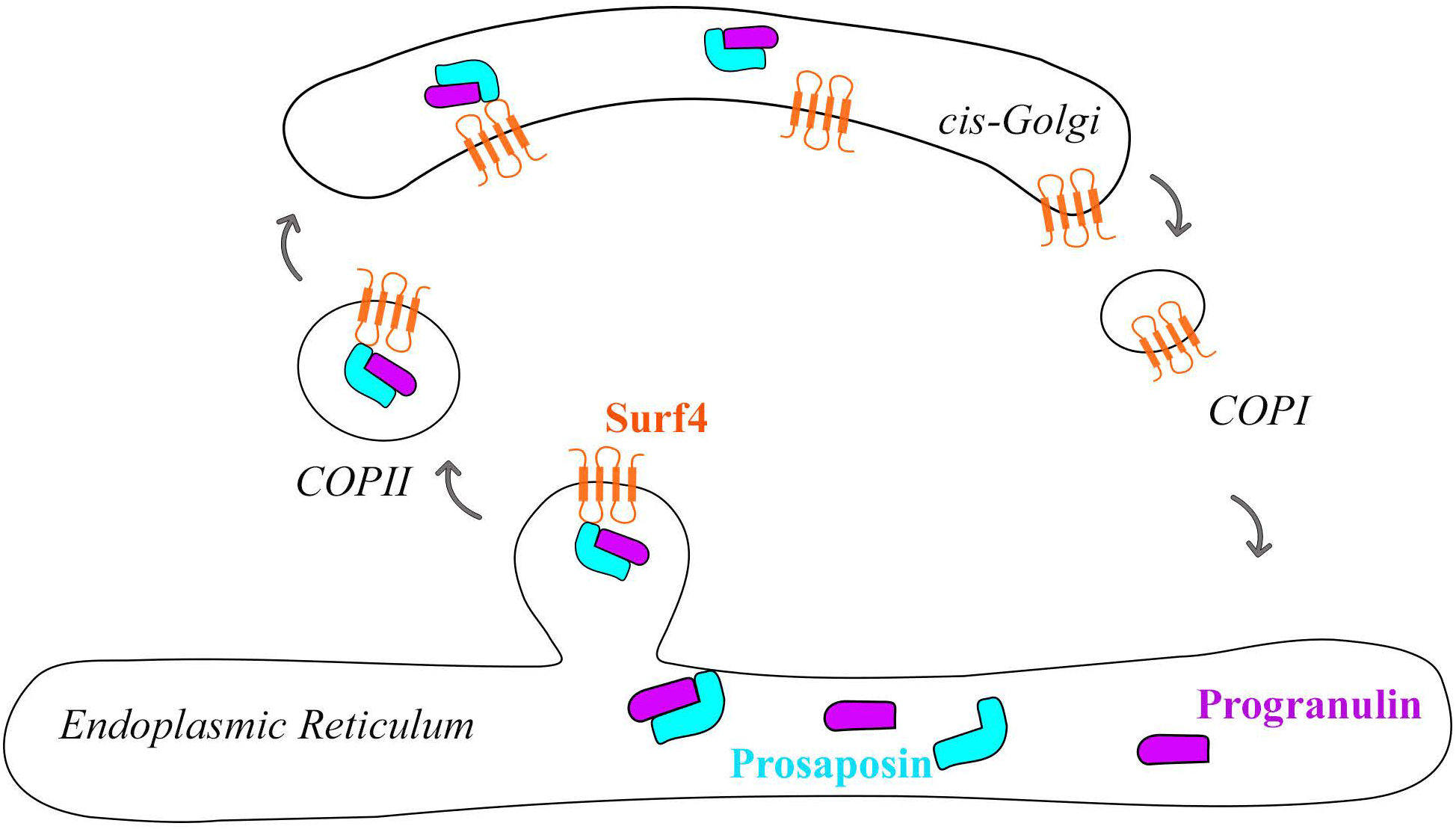
Model for Surf4-dependent ER exit of prosaposin-progranulin complex. Prosaposin forms a protein complex with progranulin in the ER and facilitates the ER to Golgi trafficking of progranulin via COPII vesicles by acting as a linker to bind to Surf4 receptor (ER-cis-Golgi outline was modified from ^56^ using Inkscape software).

Our discovery of a role for Surf4 in promoting the ER exit of progranulin and prosaposin parallels recent studies that have identified roles for CLN6 and CLN8 proteins in the sorting of other lysosomal proteins at the ER ^23,24^. Like mutations in the *GRN* gene, *CLN6* and *CLN8* mutations also cause the lysosome storage disease known as neuronal ceroid lipofuscinosis ^36,37^. As progranulin does not depend on the CLN6/8-dependent export mechanism (Supp. Fig. S3), the Surf4-dependent trafficking of progranulin and prosaposin from the ER represents a distinct mechanism that enhances ER export of progranulin and prosaposin beyond what can be achieved via bulk flow. The functionally important relationship between prosaposin and Surf4 raises new questions about how prosaposin is selectively recognized by Surf4. Although Surf4 was not previously demonstrated to sort any other mammalian lysosomal cargos, the budding yeast Surf4 homolog known as Erv29 supports trafficking of carboxypeptidase Y (CPY) to the vacuole (equivalent to the mammalian lysosome) ^38^. Our discoveries related to progranulin and prosaposin might also set a paradigm for the ER-to-Golgi trafficking of other mammalian proteins of the lysosomal lumen.

Although bulk flow has been proposed as a major mechanism to explain how lumenal proteins exit the ER, one reason for receptors to promote the efflux of specific cargos from the ER is to prioritize the export of cargos with the propensity to form aggregates ^21^. Such a mechanism has been characterized for a subset of extracellular matrix proteins that depend on Surf4 ^39^. Consistent with a need for prioritizing the ER exit of progranulin and prosaposin, we observed that they both accumulated in the ER of Surf4 KO cells. Although the basis for these accumulations remains uncertain, one possible factor is that the very cysteine rich nature of progranulin and prosaposin could create a vulnerability for the formation of non-specific disulfide bonds if their abundances exceed the capacity for protein disulfide isomerases to handle them. Although defining direct functions of progranulin and the granulins that derive from it within lysosomes remains a work in progress, the coordinated trafficking of progranulin and prosaposin suggests a possible functional relationship between these two proteins and raises questions about the evolutionary origin of their interaction. With respect to a possible functional relationship, the 4 saposin domains that are liberated from prosaposin by lysosomal proteases promote the breakdown of several species of lipids by promoting their extraction from membranes and presentation to specific hydrolases ^15,16^. For example, saposin C presents glucosylsphingosine and glucosylceramide to glucocerebrosidase (GCase, encoded by the *GBA1* gene). Mutations in *GBA1* cause Gaucher’s disease (homozygous) and confer significant Parkinson’s disease risk (heterozygous) ^40^. Therefore, the Surf4-dependence of prosaposin trafficking may have relevance for understanding these diseases and for therapies based on promoting the activity of GCase. Although much remains to be defined about the roles of individual granulins in the regulation of lysosomal hydrolases, recent studies have proposed that they regulate the activity of selected cathepsin proteases ^41–43^. Another interesting protease connection linked to progranulin sorting mechanisms comes from plants where proteins with individual granulins are fused to papain-like cysteine proteases ^44,45^. Thus, while the strong dependence of progranulin on prosaposin for its trafficking suggests the possibility of functional relationships between these proteins within lysosomes, a definitive test of this idea first requires a better understanding of the functions of progranulin and the individual granulin domains that are liberated from it within lysosomes.

In summary, we have established that ER exit and lysosomal availability of progranulin depends on an interaction with prosaposin that begins soon after their co-translational insertion into the ER (Figure 8). The complex of progranulin and prosaposin in turn depends on an interaction between prosaposin and Surf4 for efficient efflux from the ER and their eventual delivery to lysosomes. Thus, although it has previously been proposed that a process of bulk flow mediates the egress of many lumenal proteins from the ER, progranulin and prosaposin are particularly dependent on Surf4 for their export. This dependence of progranulin and prosaposin on Surf4 parallels (but is distinct from) other recent reports of lysosomal hydrolases relying on CLN6 and CLN8 proteins for their ER export ^23,24^. Beyond defining a critical step in the intracellular traffic of progranulin and prosaposin, this study sheds light on how the sorting of cargoes is coordinated between organelles and has implications for therapeutic strategies designed to increase progranulin levels as a therapy for frontotemporal dementia arising from its deficiency.

## Methods

### Plasmids and DNA Cloning

RUSH constructs were cloned using Gibson assembly (NEB #E2611S). EGFP-GalT was a gift from Jennifer Lippincott-Schwartz (Addgene plasmid # 11929; http://n2t.net/addgene:11929; RRID:Addgene_11929) ^46^. Str-KDEL_SBP-mCherry-GPI was a gift from Franck Perez (Addgene plasmid # 65295; http://n2t.net/addgene:65295; RRID:Addgene_65295) ^20^. For cloning of RUSH-Progranulin (Str-KDEL_SBP-mCherry-Progranulin), human progranulin was PCR amplified from AAVS1_Puro_PGK_mCherry-Progranulin ^47^ and the plasmid backbone was amplified from Str-KDEL_SBP-mCherry-GPI (Addgene # 65295) which supports co-expression (via an IRES linker) of the Str-KDEL and the SBP-fusion protein of interest. The IL-2 signal sequence that was present in the original mCherry-progranulin plasmid was later replaced with progranulin signal sequence by site-directed mutagenesis (NEB Q5 site directed mutagenesis kit). For cloning of Prosaposin-RUSH (Str-KDEL_Prosaposin-SBP-mCherry), prosaposin was amplified from human prosaposin cDNA (pCMV6-XL5-PSAP, Origene, # SC118405), SBP-mCherry and the Str-KDEL_SBP-mCherry-GPI (Addgene # 65295) was amplified in parallel. For cloning of Progranulin-RUSH (Str-KDEL_Progranulin-SBP-mCherry), progranulin was amplified from RUSH-Progranulin, SBP-mCherry insert and vector were amplified from Str-KDEL_SBP-mCherry-GPI (Addgene # 65295). The eGFP-Surf4 plasmid (pLV[Exp]-Puro-EF1A>eGFP-3xGS-hSurf4) was designed to fuse eGFP to the N-terminus of human Surf4 (NM_033161.4) in a mammalian lentiviral gene expression vector (VectorBuilder). eGFP-Surf4 Δmotif and eGFP-Surf4 FF→AA mutant plasmids were generated from the eGFP-Surf4 plasmid by site-directed mutagenesis. The CRISPR px459 plasmid (Addgene #62988) was kindly provided by Feng Zhang (MIT). For expression of mCherry-progranulin from the AAVS1 safe harbor locus, the hCas9 and gRNA_AAVS1-T2 plasmids were gifts from George Church (Addgene plasmid # 41815; http://n2t.net/addgene:41815; RRID:Addgene_41815 and Addgene plasmid # 41818; http://n2t.net/addgene:41818; RRID:Addgene_41818) ^48^. pcDNA3 was from Invitrogen. Oligonucleotide primer sequences are summarized in Supp. table 1.

### Cell culture

HeLa cells and mouse embryonic fibroblasts (MEFs) were maintained in DMEM supplemented with 10% fetal bovine serum (Gibco) and 1% penicillin - streptomycin mix in a humidified 37°C incubator with 5% CO_2_.

HeLa cells were seeded at 100,000 cells in 2 ml media per well in a six well dish, and transfected with 333 ng each of hCas9 (Addgene plasmid # 41815); ^48^, gRNA_AAVS1-T2 (Addgene plasmid # 41818); ^48^, and AAVS1_Puro_PGK_mCherry-Progranulin ^47^ plasmids added to the 100µl of Opti-MEM and 3µl Fugene 6 transfection mix pre-incubated for 20min. 48 hours after transfection, the transfected cells were selected with puromycin (2µg/ml) for two weeks to isolate the surviving cells that are stably expressing mCherry-progranulin.

An immortalized line of CLN6 mutant MEFs (as well as a wildtype littermate control) was generated by serial passaging of primary MEFs from the *nclf* line of spontaneous mutant mice that was obtained from The Jackson Laboratory ^36,49^.

### Generation of knockout cells by CRISPR/Cas9 gene editing

For the generation of Surf4 KO HeLa cells, sgRNA targeting exon 3 of Surf4 gene (5’-GGTGTTGAGCAGGAACTTCG -3’) was cloned into mammalian expression vector pX459 and confirmed by DNA sequencing ^50^. 0.4ug of plasmid was added to 21 µl Opti-MEM reduced serum medium (Gibco) and 1.2 µl Fugene 6 (Promega) transfection mix and incubated for 15min at room temperature before adding to HeLa cells plated in one well of a 24 well plate. The next day, the transfected cells were selected with puromycin (2µg/ml) for 48 hours. The cells were plated at low density to isolate clones derived from single cells. To identify indels near the sgRNA target region, genomic DNA was isolated (Quick Extract DNA extraction solution, Lucigen), amplified by PCR, followed by blunt-end ligation of the amplified DNA into TOPO cloning vector (Zero Blunt TOPO PCR cloning kit, Invitrogen) and transformed into TOP10 *E*.*coli*. 18 colonies were screened by isolating plasmids and indel mutations were confirmed by sequencing using the M13 forward sequencing primer. Generation of progranulin KO HeLa cells by CRISPR RNP electroporation was described previously ^47^ and indel mutations (1 bp insertion) were confirmed by DNA sequencing. DNA sequencing was performed by Keck DNA sequencing facility, Yale School of Medicine.

### siRNA transfection

For siRNA transfection, 7.5µl of 20µM siRNA was added to 500µl Opti-MEM reduced serum medium (Gibco) followed by the addition of 5µl Lipofectamine RNAi max reagent (Invitrogen) and gently mixed. After incubating the mix for 15min at room temperature, HeLa cells plated at 100,000 cells in 2ml media per well in a six-well dish were transfected for 48 hours before further analysis by immunostaining and immunoblotting. Prosaposin siRNA (5’-GGCCGACAUAUGCAAGAACUAUATC-3’) and negative control siRNA (5′-CGUUAAUCGCGUAUAAUACGCGUAT-3′) were purchased from IDT.

### RUSH live cell imaging and quantification

HeLa cells were plated at 100,000cells in 2 ml media onto 35mm MatTek glass bottom dishes and transfected with 0.9µg RUSH plasmid and 0.1µg eGFP-GalT or LAMP1-GFP using 100µl Opti-MEM and 3µl Fugene 6 transfection reagent (Promega) one day before imaging. Live cell imaging was performed using confocal laser scanning microscope equipped with an airyscan detector (Zeiss LSM880) in an environment-controlled chamber set at 37°C and 5% CO_2._ Time lapse images were acquired at 3min intervals for one hour after the addition of Biotin (40µM) to the medium using a Plan Apochromat 63x objective (N.A 1.4). Images were quantified using ImageJ ^51^. To quantify the Golgi localization of Prosaposin-RUSH and Progranulin-RUSH proteins, the eGFP-GalT labeled Golgi region was masked by setting up an intensity threshold. In this masked region, the fluorescence intensities of Cherry-tagged RUSH proteins were quantified at 3min time intervals after the addition of biotin and were normalized to images taken before the addition of biotin.

### Transient transfection of plasmids

For transient transfection of eGFP-Surf4 plasmids, cells were plated on 12-mm No.1 glass coverslips at 20,000 cells in 0.5ml media per well in a 24-well dish and were transfected with 0.25µg plasmid using 25ul Opti-MEM and 0.75ul Fugene 6 transfection mix that was pre-incubated for 20min. 48 hours after transfection, cells were used for immunostaining.

### Immunostaining, Immunofluorescence, and image quantification

For immunostaining, cells seeded on 12-mm No.1 glass coverslips (Carolina Biological supply) were fixed for 30 min at room temperature with 8% PFA in 0.1M sodium phosphate buffer (pH 7.2) that was added 1:1 to the cell culture media. Cells were then rinsed once with PBS and permeabilized for 15 min with 0.1% saponin prepared in PBS followed by incubation with blocking buffer (3% BSA and 0.1% saponin in PBS) for 20min. Cells were incubated with primary antibodies overnight at 4°C and secondary antibodies for 30min at room temperature. Primary and secondary antibodies were diluted in blocking buffer. The coverslips were washed three times with PBS containing 0.1% saponin after incubation with antibodies and mounted on glass slides using prolong gold mounting medium (Invitrogen, Carlsbad, CA).

The primary antibodies used for the immunostaining were mouse anti-LAMP1 (DSHB; catalog no # H4A3), rabbit anti-saposin C (Santa Cruz; clone H-81 catalog # SC-32587), goat anti-progranulin (R&D systems; catalog no # AF2420), rabbit anti-calnexin (Cell Signaling technology; Catalog no #C5C9), mouse anti-GM130 (BD Transduction Laboratories; Catalog no # 610822), goat anti-cathepsin D (R&D systems; Catalog no # AF1014), sheep anti-mouse progranulin (R&D systems; Catalog no # AF2557), mouse anti-LAMP1 (DSHB; Catalog no # 1D4B).

Alexa Fluor conjugated secondary antibodies used for the immunostaining were purchased from Invitrogen Life Technologies. Images were acquired using confocal laser scanning microscope equipped with an airyscan detector (Zeiss LSM880) using a Plan Apochromat 63x objective (N.A 1.4). Images were quantified for the colocalization of progranulin and prosaposin with LAMP1 per cell using CellProfiler ^52^. The Mander’s colocalization coefficient was measured using the measure correlation module by setting the Otsu automatic intensity thresholding for puncta of size above 0.294µm.

### Immunoprecipitation and Immunoblotting

To prepare protein lysates, cells were washed twice with cold PBS and scraped off the dish using cold lysis buffer (50mM Tris pH 7.4, 150mM NaCl, 1mM EDTA and 1%Triton-X100) along with protease and phosphatase inhibitors (Complete, Mini-EDTA free, PhosSTOP) and centrifuged at 14,000 rpm for 6 min at 4°C to collect the supernatant leaving behind the insoluble pellet. For immunoprecipitation, protein lysates were added to RFP-Trap beads (Chromotek) and incubated for one hour at 4°C on a rotating shaker. Beads were then washed four times with lysis buffer and eluted in 2x Laemmli buffer containing 1% 2-mercaptoethanol (Sigma) by heating at 95°C for 3min.

For immunoprecipitation of eGFP-Surf4 protein complexes, HeLa cells plated on 10cm dishes were transiently transfected with pcDNA3, Prosaposin-RUSH, eGFP-Surf4, and Prosaposin-RUSH+eGFP-Surf4 plasmids. Equal amounts of PSAP-RUSH and eGFP-Surf4 plasmids were used per dish. In dishes where PSAP-RUSH or eGFP-Surf4 were transfected alone, empty pcDNA3 was added to make up the total amount of plasmid DNA to 7µg per dish. 7µg plasmid was added to 700µl Opti-MEM and 21µl Fugene 6 transfection mix and incubated for 20min before adding to the cells. One day after transfection, cells were rinsed twice with PBS and protein crosslinking reaction was performed by incubating with freshly prepared 0.5mM DSP (ThermoFisher Scientific) in PBS for 30min at room temperature followed by quenching the reaction with 20mM Tris pH 7.4 for 10 min. Cells were rinsed twice with cold PBS and lysed in cold lysis buffer (50mM Tris pH 7.4, 150mM NaCl, 1% NP-40) along with protease and phosphatase inhibitors (Complete Mini-EDTA free, PhosSTOP), and centrifuged at 14,000 rpm for 6 min at 4°C to collect the supernatant leaving behind the insoluble pellet. Immunoprecipitation of GFP-Surf4 protein complexes was performed using GFP-Trap (Chromotek) similar to RFP-Trap IP experiment described above. To detect GFP-Surf4 on immunoblots, protein samples were denatured in Laemmli buffer for 10 min at room temperature before running on SDS-PAGE gels.

Proteins were separated on SDS-PAGE gels (BioRad; 4–15% gradient Mini-PROTEAN TGX precast polyacrylamide gels) and transferred to nitrocellulose membranes using wet blot transfer system (BioRad). Membranes were blocked with 5% non-fat dry milk prepared in TBST (Tris Buffered Saline containing 0.1% Tween-20) for one hour at room temperature followed by incubation with primary antibodies diluted in 5% BSA prepared in TBST at 4°C overnight. The next day, membranes were incubated with the corresponding HRP-conjugated secondary antibodies diluted in 5% non-fat dry milk in TBST for one hour at room temperature. Membranes were washed three times with TBST after antibody incubations. Membranes were developed using ECL chemiluminescence substrate for HRP to detect protein signal (ThermoFisher Scientific) using a Versadoc imaging system (BioRad). Immunoblots were quantified using ImageJ.

The primary antibodies used for the immunoblotting were rabbit anti-saposin C (Santa Cruz; clone H-81 catalog # SC-32587), goat anti-progranulin (R&D systems; catalog no # AF2420), rabbit anti-calnexin (Cell Signaling technology; Catalog no #C5C9), mouse anti-vinculin (Sigma; Catalog no #v4505; Clone vin -11-5), Rabbit anti-RFP (Rockland Immunochemicals; Catalog no # 600-401-379) and anti-GFP-HRP (Rockland Immunochemicals; Catalog no #600-103-215).

The secondary antibodies used were anti-biotin-HRP (Cell Signaling technology; Catalog no #7075P5), anti-rabbit-HRP (Cell Signaling technology; Catalog no #7074s), anti-mouse-HRP (Cell Signaling technology; Catalog no #7076s), anti-goat-HRP (Invitrogen; Catalog no #31402).

### Endoglycosidase H assays

Protein lysates were prepared in a lysis buffer containing 1%Triton-X100 as described above. To prepare media samples, cells were rinsed twice with warm PBS to replace DMEM with serum-free DMEM one day before collection of media. 2.5ml of media was concentrated to 50µl using centrifugal filters (Amicon Ultra-4 10kDa cutoff filters) by centrifugation at 4,000 rpm at

4°C for 30 min. 20µg of protein lysates and media samples were incubated with glycoprotein denaturation buffer at 95°C for 10 min and treated with 0.3µl Endo H (NEB; Catalog no #P0702) enzyme in glyco-buffer 3 for one hour at 37°C. After Endo H digestion, samples were denatured in Laemmli buffer by heating at 95°C for 5 min and resolved on SDS-PAGE gel for immunoblotting as above. To perform Endo H assays on immunoprecipitated samples, beads were eluted in glycoprotein denaturation buffer at 95°C for 10 min followed by Endo H treatment as described above.

### Protein sequence analysis

Human Surf4 protein sequence was analyzed for amino acid sequence conservation across species using MultAlin algorithm ^53^. To identify protein motifs, human Surf4 protein sequence was scanned against the PROSITE collection of Motifs using ScanProsite (https://prosite.expasy.org/scanprosite/) ^54,55^.

### Statistical analysis

All statistical tests were performed using Prism (GraphPad). All data were represented as mean ± SEM. Kolmogorov-Smirnov (KS) normality tests were performed to check the Gaussian distribution of data points. Statistical tests performed were indicated along with *p* values in figure legends.

## Acknowledgements

We are grateful for microscopy resources provided by the Yale Program in Cellular Neuroscience, Neurodegeneration and Repair imaging facility. This research was supported by grants from the Bluefield Project to Cure FTD (SD and SMF), The Parkinson’s Disease Foundation (SMF) and the NIH (AG062210 to SMF). The authors do not have any conflicts to declare.

## Author Contributions

SD and SMF designed experiments. SD performed all experiments. SD and SMF prepared the manuscript.

## References

1. Baker, M. et al. Mutations in progranulin cause tau-negative frontotemporal dementia linked to chromosome 17. Nature (2006). doi:10.1038/nature05016

2. Cruts, M. et al. Null mutations in progranulin cause ubiquitin-positive frontotemporal dementia linked to chromosome 17q21. Nature (2006). doi:10.1038/nature05017

3. Gass, J. et al. Mutations in progranulin are a major cause of ubiquitin-positive frontotemporal lobar degeneration. Hum. Mol. Genet. (2006). doi:10.1093/hmg/ddl241

4. Smith, K. R. et al. Strikingly different clinicopathological phenotypes determined by progranulin-mutation dosage. Am. J. Hum. Genet. (2012). doi:10.1016/j.ajhg.2012.04.021

5. Tanaka, Y., Chambers, J. K., Matsuwaki, T., Yamanouchi, K. & Nishihara, M. Possible involvement of lysosomal dysfunction in pathological changes of the brain in aged progranulin-deficient mice. Acta Neuropathol. Commun. 2, 78 (2014).

6. Cenik, B., Sephton, C. F., Cenik, B. K., Herz, J. & Yu, G. Progranulin: A proteolytically processed protein at the crossroads of inflammation and neurodegeneration. Journal of Biological Chemistry (2012). doi:10.1074/jbc.R112.399170

7. Holler, C. J., Taylor, G., Deng, Q. & Kukar, T. Intracellular proteolysis of progranulin generates stable, lysosomal granulins that are haploinsufficient in patients with frontotemporal dementia caused by GRN mutations. eNeuro (2017). doi:10.1523/ENEURO.0100-17.2017

8. Kao, A. W., McKay, A., Singh, P. P., Brunet, A. & Huang, E. J. Progranulin, lysosomal regulation and neurodegenerative disease. Nat. Rev. Neurosci. 18, 325–333 (2017).

9. Lee, C. W. et al. The lysosomal protein cathepsin L is a progranulin protease. Mol. Neurodegener. 12, 55 (2017).

10. Petkau, T. L. & Leavitt, B. R. Progranulin in neurodegenerative disease. Trends Neurosci. 37, 388–98 (2014).

11. Zhou, X. et al. Lysosomal processing of progranulin. Mol. Neurodegener. (2017). doi:10.1186/s13024-017-0205-9

12. Hu, F. et al. Sortilin-mediated endocytosis determines levels of the frontotemporal dementia protein, progranulin. Neuron 68, 654–67 (2010).

13. Nicholson, A. M. et al. Prosaposin is a regulator of progranulin levels and oligomerization. Nat. Commun. (2016). doi:10.1038/ncomms11992

14. Zhou, X. et al. Prosaposin facilitates sortilin-independent lysosomal trafficking of progranulin. J. Cell Biol. (2015). doi:10.1083/jcb.201502029

15. Kishimoto, Y., Hiraiwa, M. & O’Brien, J. S. Saposins: structure, function, distribution, and molecular genetics. J. Lipid Res. 33, 1255–67 (1992).

16. Meyer, R. C., Giddens, M. M., Coleman, B. M. & Hall, R. A. The protective role of prosaposin and its receptors in the nervous system. Brain Research (2014). doi:10.1016/j.brainres.2014.08.022

17. Zhou, X., Kukar, T. & Rademakers, R. Lysosomal Dysfunction and Other Pathomechanisms in FTLD: Evidence from Progranulin Genetics and Biology. Adv. Exp. Med. Biol. (2021). doi:10.1007/978-3-030-51140-1_14

18. Carrasquillo, M. M. et al. Genome-wide screen identifies rs646776 near sortilin as a regulator of progranulin levels in human plasma. Am. J. Hum. Genet. 87, 890–897 (2010).

19. Freeze, H. H. & Kranz, C. Endoglycosidase and glycoamidase release of n-linked glycans. Current Protocols in Molecular Biology (2010). doi:10.1002/0471140864.mb1713as48

20. Boncompain, G. et al. Synchronization of secretory protein traffic in populations of cells. Nat. Methods (2012). doi:10.1038/nmeth.1928

21. Barlowe, C. & Helenius, A. Cargo Capture and Bulk Flow in the Early Secretory Pathway. Annu. Rev. Cell Dev. Biol. (2016). doi:10.1146/annurev-cellbio-111315-125016

22. Gomez-Navarro, N. & Miller, E. Protein sorting at the ER-Golgi interface. J. Cell Biol. (2016). doi:10.1083/jcb.201610031

23. Bajaj, L. et al. A CLN6-CLN8 complex recruits lysosomal enzymes at the ER for Golgi transfer. J. Clin. Invest. (2020). doi:10.1172/JCI130955

24. di Ronza, A. et al. CLN8 is an endoplasmic reticulum cargo receptor that regulates lysosome biogenesis. Nat. Cell Biol. 20, 1370–1377 (2018).

25. Emmer, B. T. et al. The cargo receptor SURF4 promotes the efficient cellular secretion of PCSK9. Elife (2018). doi:10.7554/eLife.38839

26. Saegusa, K., Sato, M., Morooka, N., Hara, T. & Sato, K. SFT-4/Surf4 control ER export of soluble cargo proteins and participate in ER exit site organization. J. Cell Biol. 217, 2073–2085 (2018).

27. Lin, Z. et al. The Endoplasmic Reticulum Cargo Receptor SURF4 Facilitates Efficient Erythropoietin Secretion. Mol. Cell. Biol. 40, 1–17 (2020).

28. Wang, X. et al. Receptor-Mediated ER Export of Lipoproteins Controls Lipid Homeostasis in Mice and Humans. Cell Metab. (2021). doi:10.1016/j.cmet.2020.10.020

29. Mitrovic, S., Ben-Tekaya, H., Koegler, E., Gruenberg, J. & Hauri, H. P. The cargo receptors Surf4, endoplasmic reticulum-Golgi intermediate compartment (ERGIC)-53, and p25 are required to maintain the architecture of ERGIC and Golgi. Mol. Biol. Cell (2008). doi:10.1091/mbc.E07-10-0989

30. Barlowe, C. Signals for COPII-dependent export from the ER: What’s the ticket out? Trends Cell Biol. 13, 295–300 (2003).

31. Foley, D. A., Sharpe, H. J. & Otte, S. Membrane topology of the endoplasmic reticulum to Golgi transport factor Erv29p. Mol. Membr. Biol. 24, 259–268 (2007).

32. Nie, C., Wang, H., Wang, R., Ginsburg, D. & Chen, X. W. Dimeric sorting code for concentrative cargo selection by the COPII coat. Proc. Natl. Acad. Sci. U. S. A. (2018). doi:10.1073/pnas.1704639115

33. Nufer, O. et al. Role of cytoplasmic C-terminal amino acids of membrane proteins in ER export. J. Cell Sci. (2002).

34. Braulke, T. & Bonifacino, J. S. Sorting of lysosomal proteins. Biochimica et Biophysica Acta - Molecular Cell Research (2009). doi:10.1016/j.bbamcr.2008.10.016

35. Ghosh, P., Dahms, N. M. & Kornfeld, S. Mannose 6-phosphate receptors: New twists in the tale. Nature Reviews Molecular Cell Biology (2003). doi:10.1038/nrm1050

36. Gao, H. et al. Mutations in a novel CLN6-encoded transmembrane protein cause variant neuronal ceroid lipofuscinosis in man and mouse. Am. J. Hum. Genet. (2002). doi:10.1086/338190

37. Ranta, S. et al. The neuronal ceroid lipofuscinoses in human EPMR and mnd mutant mice are associated with mutations in CLN8. Nat. Genet. (1999). doi:10.1038/13868

38. Belden, W. J. & Barlowe, C. Role of Erv29p in collecting soluble secretory proteins into ER-derived transport vesicles. Science (80-.). (2001). doi:10.1126/science.1065224

39. Yin, Y. et al. Surf4 (Erv29p) binds amino-terminal tripeptide motifs of soluble cargo proteins with different affinities, enabling prioritization of their exit from the endoplasmic reticulum. PLoS Biol. (2018). doi:10.1371/journal.pbio.2005140

40. Avenali, M., Blandini, F. & Cerri, S. Glucocerebrosidase Defects as a Major Risk Factor for Parkinson’s Disease. Frontiers in Aging Neuroscience (2020). doi:10.3389/fnagi.2020.00097

41. Beel, S. et al. Progranulin functions as a cathepsin D chaperone to stimulate axonal outgrowth in vivo. Hum. Mol. Genet. 26, 2850–2863 (2017).

42. Valdez, C. et al. Progranulin-mediated deficiency of cathepsin D results in FTD and NCL-like phenotypes in neurons derived from FTD patients. Hum. Mol. Genet. (2017). doi:10.1093/hmg/ddx364

43. Butler, V. J. et al. Multi-Granulin Domain Peptides Bind to Pro-Cathepsin D and Stimulate Its Enzymatic Activity More Effectively Than Progranulin in Vitro. Biochemistry (2019). doi:10.1021/acs.biochem.9b00275

44. Avrova, A. O. et al. A cysteine protease gene is expressed early in resistant potato interactions with Phytophthora infestans. Mol. Plant-Microbe Interact. (1999). doi:10.1094/MPMI.1999.12.12.1114

45. Gu, C. et al. Post-translational regulation and trafficking of the granulin-containing protease rd21 of arabidopsis thaliana. PLoS One (2012). doi:10.1371/journal.pone.0032422

46. Cole, N. B. et al. Diffusional mobility of Golgi proteins in membranes of living cells. Science (80-.). (1996). doi:10.1126/science.273.5276.797

47. Nguyen, A. D. et al. Murine knockin model for progranulin-deficient frontotemporal dementia with nonsensemediated mRNA decay. Proc. Natl. Acad. Sci. U. S. A. (2018). doi:10.1073/pnas.1722344115

48. Mali, P. et al. RNA-guided human genome engineering via Cas9. Science (80-.). (2013). doi:10.1126/science.1232033

49. Todaro, G. J. & Green, H. Quantitative studies of the growth of mouse embryo cells in culture and their development into established lines. J. Cell Biol. (1963). doi:10.1083/jcb.17.2.299

50. Ran, F. A. et al. Genome engineering using the CRISPR-Cas9 system. Nat. Protoc. (2013). doi:10.1038/nprot.2013.143

51. Schneider, C. A., Rasband, W. S. & Eliceiri, K. W. NIH Image to ImageJ: 25 years of image analysis. Nature Methods (2012). doi:10.1038/nmeth.2089

52. Carpenter, A. E. et al. CellProfiler: Image analysis software for identifying and quantifying cell phenotypes. Genome Biol. (2006). doi:10.1186/gb-2006-7-10-r100

53. Corpet, F. Multiple sequence alignment with hierarchical clustering. Nucleic Acids Res. (1988). doi:10.1093/nar/16.22.10881

54. de Castro, E. et al. ScanProsite: Detection of PROSITE signature matches and ProRule-associated functional and structural residues in proteins. Nucleic Acids Res. (2006). doi:10.1093/nar/gkl124

55. Sigrist, C. J. A. et al. New and continuing developments at PROSITE. Nucleic Acids Res. (2013). doi:10.1093/nar/gks1067

56. Bykov, Y. S. et al. The structure of the COPI coat determined within the cell. Elife (2017). doi:10.7554/eLife.32493

