## Supplemental Material for "Surf4 Promotes Endoplasmic Reticulum Exit of the Lysosomal Prosaposin-Progranulin Complex"

Devireddy, S.[1,2,3], and Ferguson, S.M. [1,2,3*]

Departments of Cell Biology[1] and Neuroscience[2], Program in Cellular Neuroscience, Neurodegeneration and Repair[3], Yale University School of Medicine, New Haven, Connecticut 06510, USA;

Running title: Surf4-dependent ER exit of progranulin-prosaposin complex

**This PDF file includes:**

Supp. Figures 1 to 5

Supp. table 1

Supp. movie 1 to 4 (Legends)

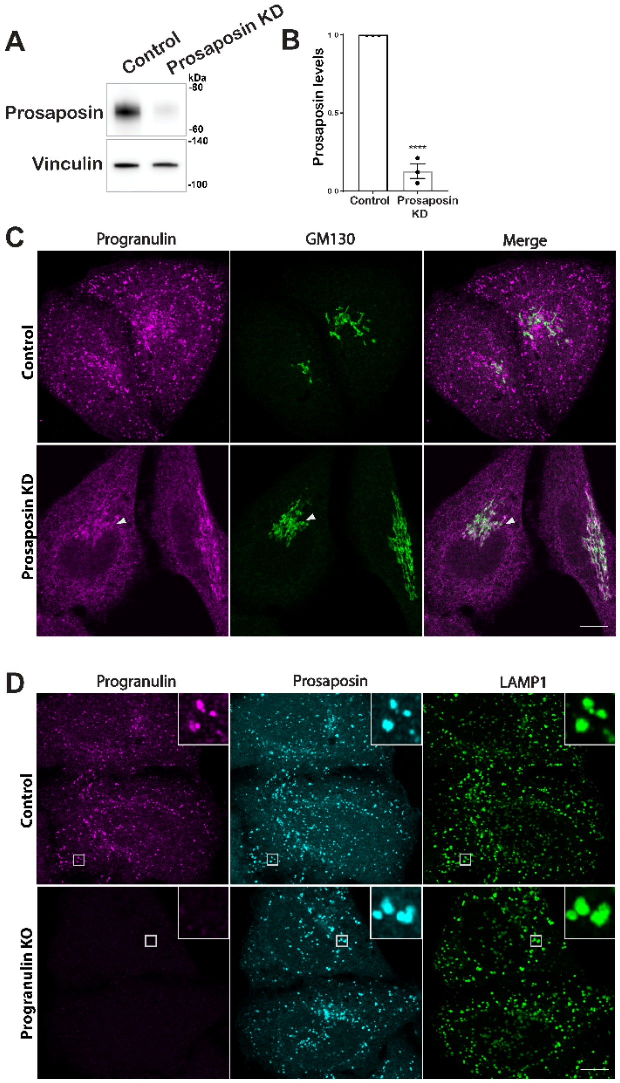

**Supp. Fig. 1: Evaluation of prosaposin and progranulin depleted cells** (A) Immunoblot evaluation of prosaposin protein levels on cells transfected with control and prosaposin siRNAs. Vinculin was used as a loading control. (B) Immunoblots were quantified by densitometry using ImageJ, and the graph shows prosaposin levels normalized to vinculin (n=3 independent experiments; mean ± SEM; Unpaired *t* test, ****$P$ value <0.0001). (C) Confocal immunofluorescence images show progranulin localization and GM130 (cis-Golgi) in control and prosaposin siRNA treated cells (scale bar, 10μm). (D) Confocal immunofluorescence images show prosaposin localization and LAMP1 (lysosomes) in control and progranulin knockout cells (scale bar, 10μm).

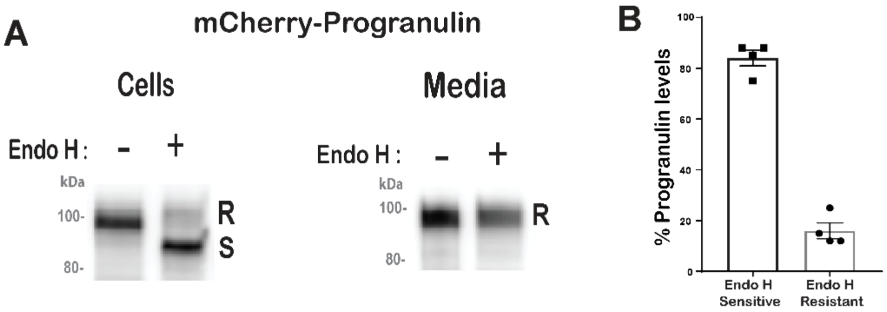

**Supp. Fig. 2: The majority of full length progranulin resides in the ER** (A) Immunoblot analysis of progranulin forms upon Endo H digestion of cell lysates and conditioned media samples collected from HeLa cells stably expressing mCherry-Progranulin. (B) Blot quantification shows the percentage of progranulin forms present in cells. n=4 experiments.

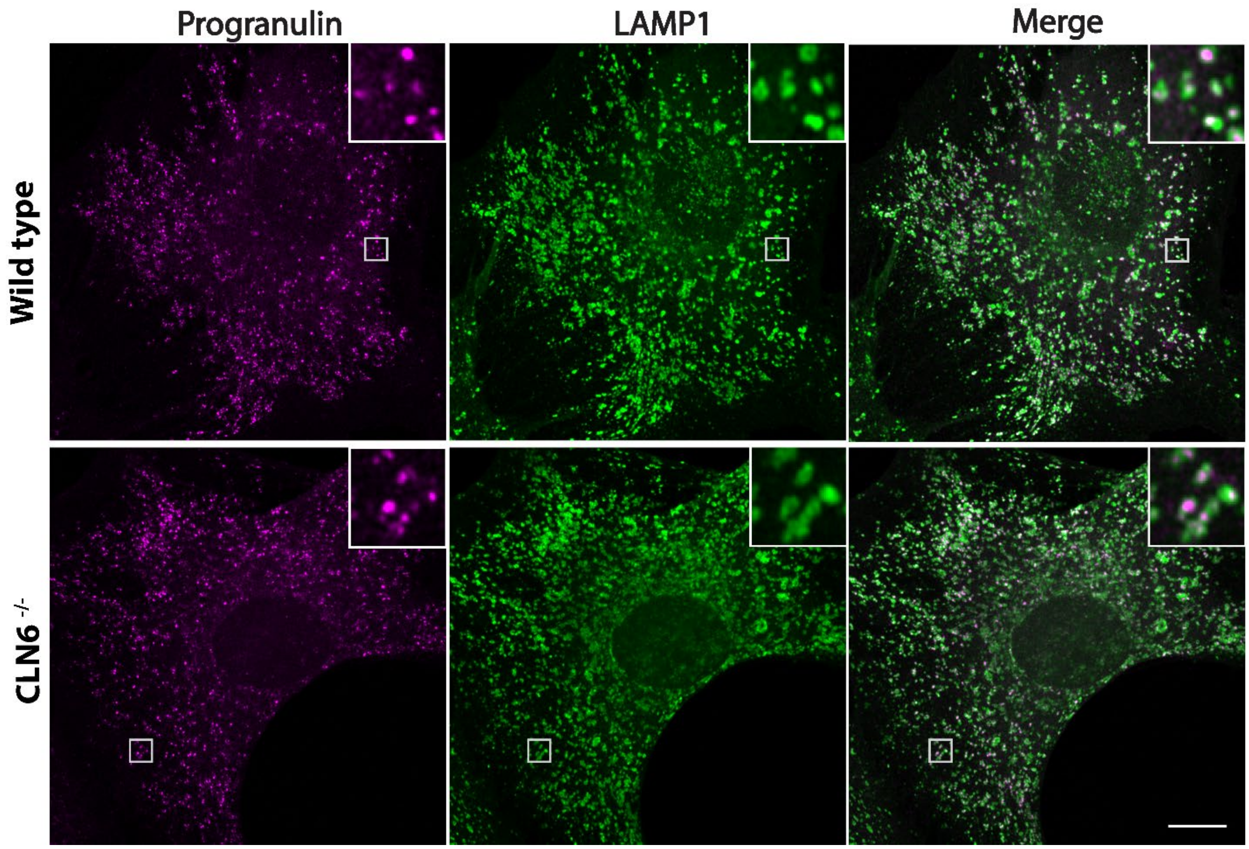

**Supp. Fig. 3: CLN6 is not required for localization of progranulin to lysosomes** Confocal immunofluorescence images show progranulin localization to LAMP1 positive late endosomes and lysosomes in embryonic fibroblasts from wildtype and CLN6 mutant mouse embryonic fibroblasts (scale bar, 10μm).

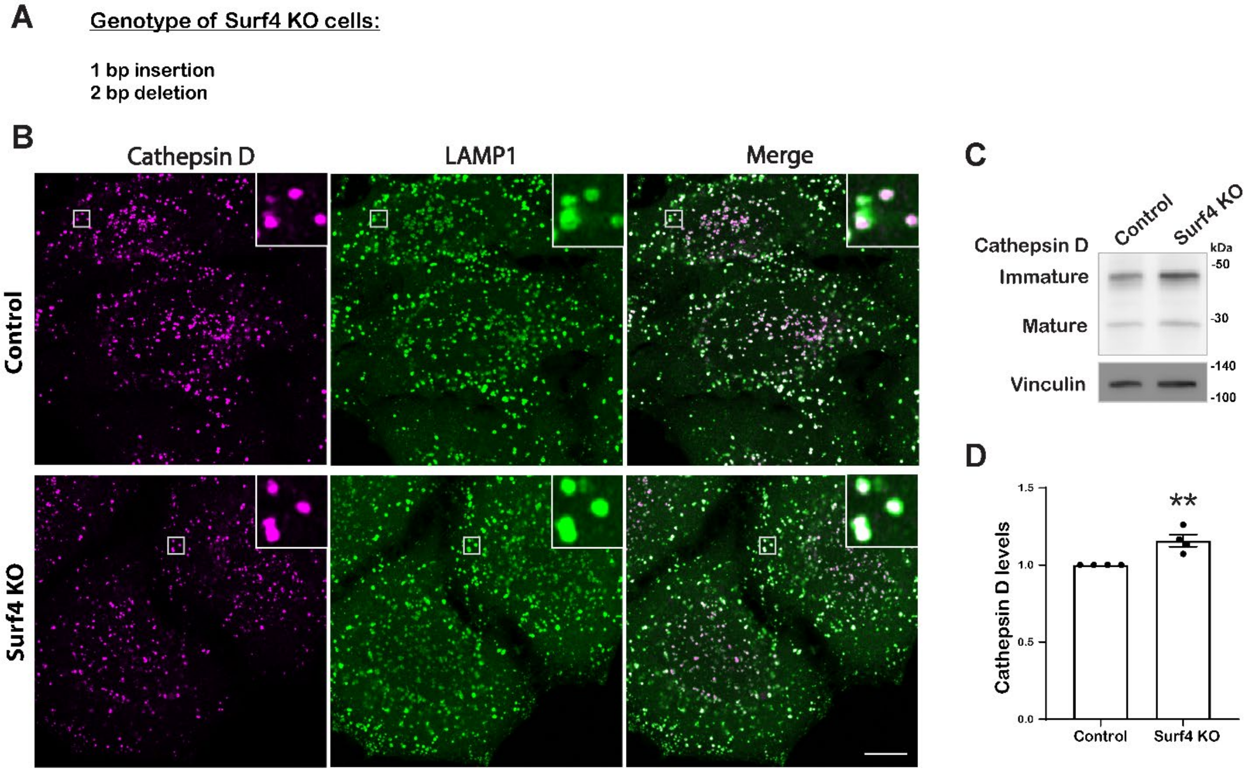

**Supp. Fig. 4: Cathepsin D trafficking is normal in the absence of Surf4** (A) Genotyping of Surf4 KO cells identified frameshift causing mutations (1 bp insertion and 2bp deletion). (B) Confocal immunofluorescence images of the localization of Cathepsin D and LAMP1 in control and Surf4 KO cells (scale bar, 10μm). (C) Immunoblot of cathepsin D protein levels in control and Surf4 KO cells. Vinculin was used as a loading control. (D) Quantification of cathepsin D levels from 4 independent experiments (mean ± SEM; unpaired *t* test; ***p*<0.01).

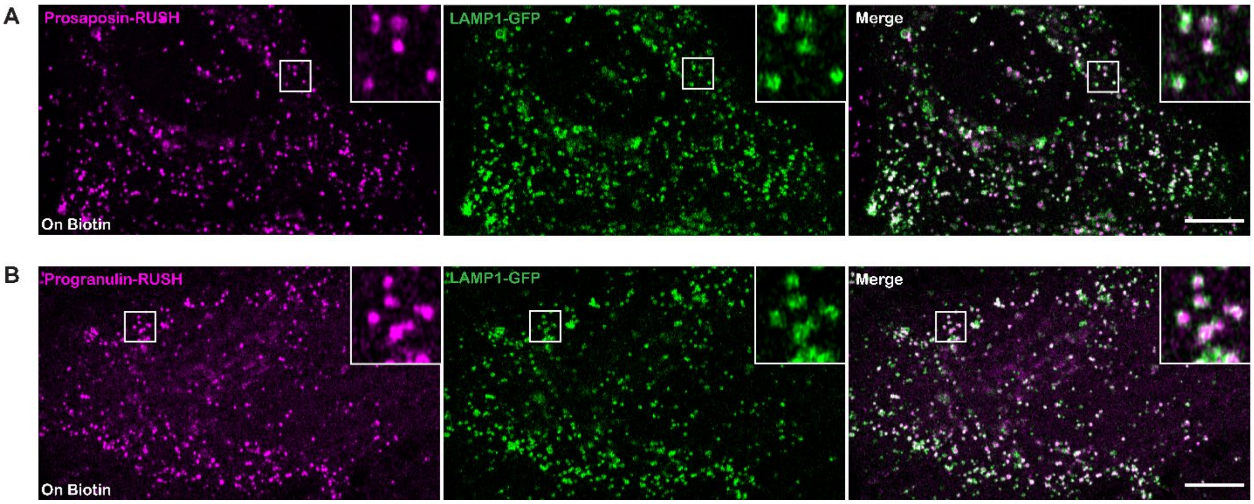

**Supp. Fig. 5: Lysosomal delivery of prosaposin-RUSH and progranulin-RUSH reporter proteins** Confocal live cell images show the localization of Prosaposin-RUSH and Progranulin-RUSH proteins to LAMP1-GFP labeled late endosomes and lysosomes in HeLa cells treated with biotin overnight (scale bars, 10μm).

### Supp. table 1: List of Oligonucleotide primer sequences

| **Plasmids** | **List of primers** | **Primers in 5` --> 3`** |
|---|---|---|
| **Surf4 gRNA in px459 vector** | gRNA Forward primer | CACCGCGAAGTTCCTGCTCAACACC |
| | gRNA Reverse primer | AAACGGTGTTGAGCAGGAACTTCGC |
| PCR primers (for genomic DNA from Surf4 KO) | Forward primer | GCAGTCCACAGTAGAGCAGT |
| | Reverse primer | ACCGAATGACCTGCCTCTAC |
| M13 sequencing primer | Forward primer | GTAAAACGACGGCCAG |
| **RUSH-Progranulin with IL2 SS construct** | | |
| Progranulin insert | Progranulin Forward primer | GGCATGGACGAGCTGTACAAGGCCGTCACGCGGTGCCCAGATGG |
| | Progranulin Reverse primer | TTAATTAATTGGCCCTCGAGGCCTCACAGCAGCTGTCTCAAGGCTG |
| Vector | Vector Forward primer | GGCCTCGAGGGCCAATTAATTAAC |
| | Vector Reverse primer | CTTGTACAGCTCGTCCATGCC |
| **RUSH-Progranulin with Progranulin SS construct** | Progranulin SS Forward primer | CTTAACAGCAGGGCTGGTGGCTGGAGACGAGAAGACCACTGGTTGGCGAGG |
| | Progranulin SS Reverse primer | GCCACCCAGCTCACCAGGGTCCACATGGCGCGCCTCCCGGGTTG |
| **Prosaposin-RUSH construct** | | |
| Prosaposin insert 1 | Prosaposin insert Forward primer | ATGTACGCCCTCTTCCTCCT |
| | Prosaposin insert Reverse primer | CGTGTCCACCTCGCCAACCAGTGGTCTTCTCGTCGACGGCGTTCCACACATGGC GTTTGC |
| SBP-mCherry insert 2 | SBP-mCherry Forward primer | GCCGTCGACGAGAAGACCAC |
| | SBP-mCherry Reverse primer | TTTCAGACAGAGAGGTTCCGCCATTTTCCAGTGGCCGGCCCTACTTGTACAGCT CGTCCA |
| Vector | Vector Forward primer | GGCCGGCCACTGGAAAATGG |
| | Vector Reverse primer | CCGCGCCCAGGAGGCTGGCCAGGAGGAAGAGGGCGTACATGGCGCGCCTCCC GGGTTGTG |
| **Progranulin-RUSH construct** | | |
| Progranulin insert 1 | Progranulin insert Forward primer | ACGCGGTGCCCAGATGGTCA |
| | Progranulin insert Reverse primer | CGTGTCCACCTCGCCAACCAGTGGTCTTCTCGTCGACGGCCAGCAGCTGTCTCA AGGCTG |

| | | |
|---|---|---|
| SBP-mCherry insert 2 | SBP-mCherry Forward primer | GCCGTCGACGAGAAGACCACTG |
| | SBP-mCherry Reverse primer | TCAGTTATCTAGAGTTAATTAATTGGCCCTCGAGGCCTCACTTGTACAGCTCGTCCATGC |
| Vector | Vector Forward primer | TGAGGCCTCGAGGGCCAATT |
| | Vector Reverse primer | AGCAGGCCACAGGGCAGAACTGACCATCTGGGCACCGCGTTCCAGCCACCAGCCCTGCTG |
| **GFP-Surf4 motif del construct** | Forward primer | CTGGTGGTGGCCCTGGGC |
| | Reverse primer | CAGGAAGTCATGCATGGGCTTGTAGAC |
| **GFP-Surf4 FF-->AA construct** | Forward primer | GAAATACGACGCTGCACAGACCATGTCGGTGATTG |
| | Reverse primer | AGGAAGTCATGCATGGGC |

### Supplementary movie Legends:

**Supp. movie 1: Prosaposin-RUSH trafficking from the ER in control cells** This video shows the trafficking of prosaposin-RUSH (magenta) from the ER in control cells beginning at 6min after biotin addition and time lapse images were acquired at 3min time intervals. eGFP-GalT labels the Golgi compartment (green). Video was played at 2 frames per second with time display in hr:min format. Scale Bar, 10μm.

**Supp. movie 2: Prosaposin-RUSH trafficking from the ER in Surf4 KO cells** This video shows the trafficking of prosaposin-RUSH (magenta) from the ER in Surf4 KO cells beginning at 6min after biotin addition and time lapse images were acquired at 3min time intervals. eGFP-GalT labels the Golgi compartment (green). Video was played at 2 frames per second with time display in hr:min format. Scale Bar, 10μm.

**Supp. movie 3: Progranulin-RUSH trafficking from the ER in control cells** This video shows the trafficking of progranulin-RUSH (magenta) from the ER in control cells beginning at 6min after biotin addition and time lapse images were acquired at 3min time intervals. eGFP-

GalT labels the Golgi compartment (green). Video was played at 2 frames per second with time display in hr:min format. Scale Bar, 10μm.

**Supp. movie 4: Progranulin-RUSH trafficking from the ER in Surf4 KO cells** This video shows the trafficking of progranulin-RUSH (magenta) from the ER in Surf4 KO cells beginning at 6min after biotin addition and time lapse images were acquired at 3min time intervals. eGFP-GalT labels the Golgi compartment (green). Video was played at 2 frames per second with time display in hr:min format. Scale Bar, 10μm.
